# Definition of the fibroblast adhesome using multiplexed proximity biotinylation

**DOI:** 10.1101/2020.01.24.918458

**Authors:** Megan R. Chastney, Craig Lawless, Jonathan D. Humphries, Stacey Warwood, Matthew C. Jones, David Knight, Claus Jorgensen, Martin J. Humphries

## Abstract

Integrin adhesion complexes (IACs) bridge the extracellular matrix to the actin cytoskeleton and transduce signals in response to both chemical and mechanical cues. The composition, interactions, stoichiometry and topological organisation of proteins within IACs are not fully understood. To address this gap, we used multiplexed proximity biotinylation (BioID) to generate an *in situ,* proximity-dependent adhesome in mouse pancreatic fibroblasts. Integration of the interactomes of 16 IAC-associated baits revealed a network of 147 proteins with 361 proximity interactions. Candidates with underappreciated roles in adhesion were identified, in addition to established IAC components. Bioinformatic analysis revealed five clusters of IAC baits that link to common groups of prey, and which therefore may represent functional modules. The five clusters, and their spatial associations, are consistent with current models of IAC interaction networks and stratification. This study provides a resource to examine proximal relationships within IACs at a global level.

## Introduction

The ability of cells to adhere to the extracellular matrix (ECM), and respond to its chemical and mechanical properties, is essential for multicellular life. This adhesion is primarily mediated by integrin receptors, which bridge the ECM to the contractile actomyosin cytoskeleton via a range of IACs ^1, 2^. In addition to providing a mechanical interface, IACs form a signalling hub from which many biochemical and biomechanical signalling pathways are transduced to guide cellular fate ^3, 4^.

The complement of adaptors, enzymes and cytoskeletal components that associate at IACs has been termed the adhesome ^5^. Extensive literature mining led to the construction of an *in silico* network of over 200 IAC-associated proteins, forming a ‘literature-curated adhesome’ ^5, 6^. The scale of the reported complexity of IACs is consistent with their substantial functional diversity. Although empirical analysis of IACs has been hampered by their lability and vicinity to the plasma membrane, advances in mass spectrometry coupled to the development of protocols to isolate IACs and adjacent material have greatly facilitated the identification of IAC-associated proteins ^7–13^. The large-scale examination of isolated IA–Cs enabled assembly of adhesome datasets, and the bioinformatic integration of seven such analyses (of fibronectin substrate-induced IACs) defined a ‘meta adhesome’ of over 2,400 proteins ^11^. This dataset was further refined to a ‘consensus adhesome’ of 60 commonly-identified components postulated to represent the core adhesion machinery. These 60 components were organised into an interaction network containing four inter-connected, hypothetical signalling modules. How closely this theoretical interaction network represents IAC organisation and protein-protein interactions (PPIs) *in situ* has yet to be experimentally defined.

Dynamic PPIs underpin the transmission of biochemical and biomechanical information across IACs, and are therefore dependent upon the organisation of adhesome components. Microscopic analyses have enabled the close examination of the ultrastructure of IACs, revealing a high degree of lateral and vertical organisation ^14–17^. For example, super-resolution light microscopy revealed the vertical stratification of a number of components within IACs at a 10-20 nm resolution, revealing three distinct, but overlapping, functional layers, albeit with some differences in the localisation of specific components ^14, 15, 18, 19^. However, the organisation of adhesome components on a larger scale has yet to be experimentally defined, and an empirical view of protein interactions within IACs is lacking.

Recently, proximity-dependent biotinylation techniques, such as BioID, have offered attractive alternatives to affinity purification approaches to examine protein proximal associations. BioID uses a mutated biotin ligase, BirA*, fused to a protein of interest (the bait), to promiscuously biotinylate proximal proteins (the prey) over the course of several hours, with an estimated labelling range of 10-15nm ^20^. As labelling occurs *in situ,* and purification takes advantage of the high affinity bond between biotin and avidin, proximity biotinylation circumvents the need to retain PPIs throughout processing. BioID has been used to probe the structure of labile and membrane-associated complexes that are difficult to study using more traditional techniques, including nuclear pore complexes, the centrosome and cell-cell contacts ^20–22^. BioID has also been used to examine the proximity interactomes of individual IAC-associated proteins, and has revealed a number of potential new adhesome candidates ^23–25^. For example, KANK2 was identified as a paxillin- and kindlin-2-proximal protein in U2OS cells, and was shown to localise to IACs ^23^. To date, however, a large-scale analysis of protein proximal networks in IACs has not been performed.

In this study, we have multiplexed BioID data from a set of 16 IAC component baits to generate a proximity-dependent adhesome. The resulting resource enables the interrogation of the proximal relationships between adhesome components, in addition to providing insights into the architecture of IACs. Bioinformatic analysis of the data revealed five clusters of bait proteins that linked to common groups of proteins with diverse, but overlapping functional roles, which may represent functional modules. The grouping of these proteins was consistent with current literature-based models of IAC interaction networks ^3, 11^. Interrogation of the topological organisation of the proximal interaction network identified a bait-prey organisation that is consistent with the reported stratified arrangement of components within IACs. A number of well-characterised adhesome components were identified among a group of 11 proteins with multiple links to a range of bait proteins, which may be part of the core adhesome machinery. This group also contained several proteins that may have underappreciated roles in adhesion regulation. This empirically-defined adhesome network provides a valuable tool to interrogate the proximity interaction networks within IACs, and to drive further hypothesis generation.

## Results and discussion

### Generation of a proximity-dependent adhesome

To maximise the capture of proximity interactions within IACs, 16 commonly-identified adhesome components were selected as BioID baits. The 16 baits represent a broad range of functions within the adhesome, and span the four putative signalling axes of the consensus adhesome (α-actinin-zyxin-VASP, talin-vinculin, FAK-paxillin, and kindlin-ILK-PINCH-kindlin; fig. 1A). The selected proteins were cloned into the pCDH lentiviral vector containing myc-BirA* with a self-cleaving blue fluorescent protein, and stably expressed in an immortalised mouse pancreatic fibroblast cell line (imPSC). Immunofluorescence microscopy confirmed colocalisation of BirA*-tagged adhesome baits and biotinylated proteins with paxillin-positive structures (supp. 1), confirming that subcellular targeting to IACs was not inhibited by the myc-BirA* tag. All baits strongly colocalised with paxillin, but in addition BirA*-PDLIM5, -palladin, -ponsin and -zyxin also stained IAC-proximal actin filaments, and BirA*-β-Pix, -GIT1, and -p130Cas staining was slightly more diffuse than other baits. Cells expressing the BirA*-only control showed no specific subcellular localisation of bait or biotinylated proteins.

**Fig. 1:**
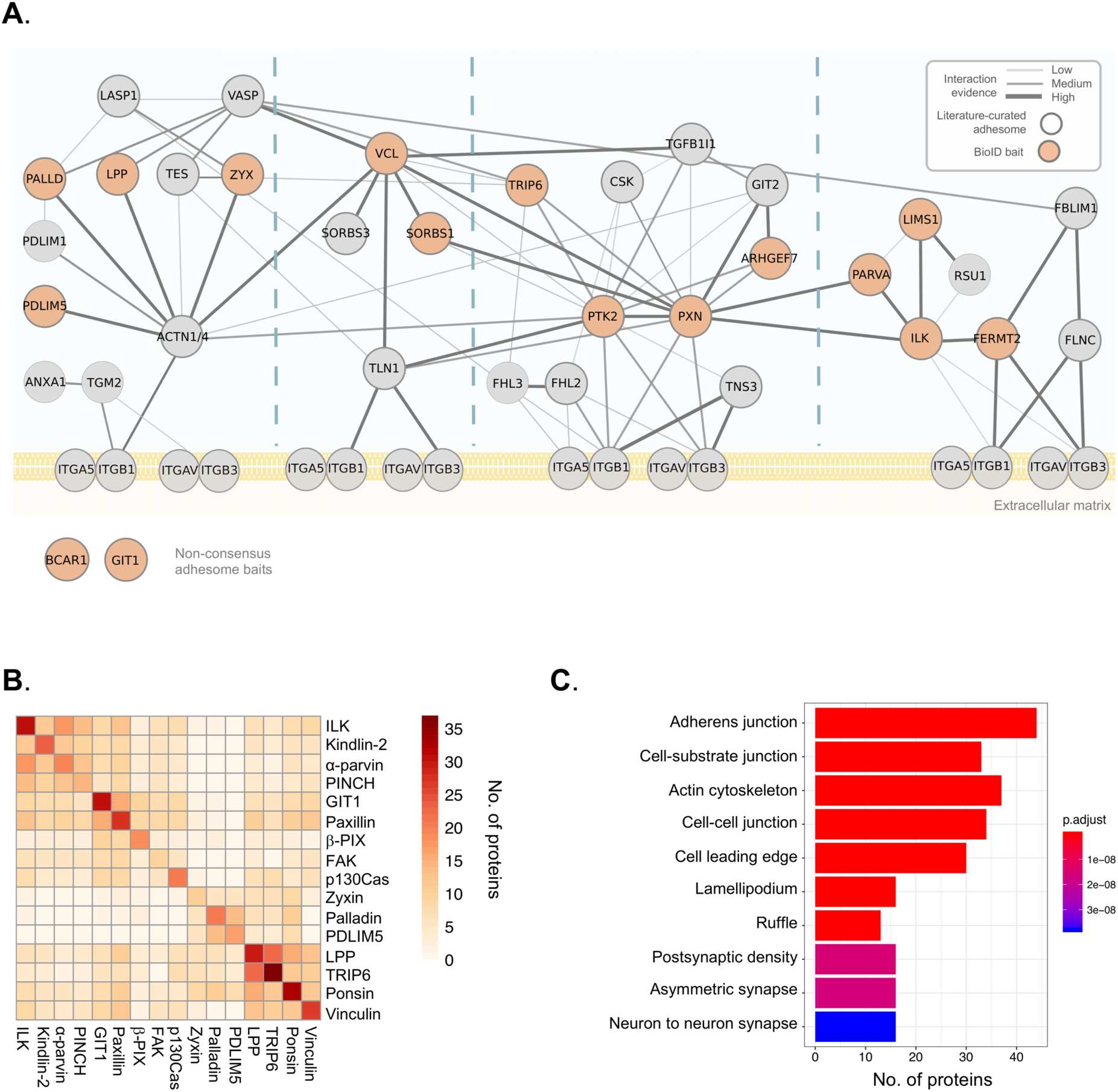
Overview of the proximity-dependent adhesome. **A.** 16 adhesome proteins were selected as BioID baits, 14 of which were present in the consensus adhesome and span the 4 putative signalling axes of the PPI network 11. p130Cas (Bcar1) and GIT1 are present in the literature-curated adhesome (and the GIT1 homolog GIT2 is in the consensus adhesome) 5. Baits are shown in orange, and edges represent evidence of PPIs. Thick grey borders indicate literature-curated adhesome proteins. Gene names are shown. Consensus adhesome components unconnected to the main network are not shown. **B.** Pairwise comparisons of proximal proteins (BFDR ≤ 0.05) identified by each BioID bait are displayed as a heatmap. Protein names not matching the gene names in (A) are: FAK, PTK; kindlin-2, FERMT2; palladin, PALLD; α-parvin, PARVA; paxillin, PXN; β-Pix, ARHGEF7; PINCH, LIMS1; ponsin, SORBS1; vinculin, VCL; zyxin, ZYX. **C.** GO enrichment analysis of the 147 proteins in the proximity-dependent adhesome. The top ten over-represented terms under the cellular component category are shown.

To determine proximal interactors of each BioID adhesome bait, label-free quantitative mass spectrometry was performed on affinity-purified biotinylated proteins from three independent experiments, and raw data were analysed by MaxQuant using ion intensity-based quantification. SAINTexpress was used to identify high-confidence bait-prey proximity interactions, with BirA* as a negative control for non-specific interactions (supp. table 1). The number of proteins predicted to be within each proximity interactome varied by bait, with 10-37 proteins predicted to be ‘true’ proximity interactors at a BFDR of ≤ 0.05. Pairwise comparisons were performed to visualise the number of proximal proteins common to each bait (fig. 1B). While some BioID baits, such as BirA*-LPP and -TRIP6, shared a large number of proteins, others showed little similarity to the majority of baits, notably BirA*-tagged zyxin, palladin and PDLIM5.

The 16 individual BioID datasets were integrated into a single network to generate a proximity-dependent adhesome. A total of 147 proximal proteins were found across all datasets, which is likely to represent a combination of core IAC proteins, IAC-associated proteins, and proteins with a proximal association with the BioID baits in more distal subcellular localisations. 361 proximity interactions were identified (excluding bait-bait interactions), the majority of which are absent from published PPI databases (see methods for details) (292; 81%). These associations may represent unknown direct interactions, indirect proximity interactions, or non-specific background interactions. Excluding BirA*-tagged bait proteins, over half of the prey identified (77) were unique to a specific bait, while eight proteins were identified by at least half of the 16 BioID baits (and may therefore represent core adhesome components) (supp. 2A).

Gene ontology (GO) analysis of the network revealed an over-representation of multiple terms related to cell-ECM adhesion, including ‘cell-substrate junction’, ‘actin cytoskeleton’ and ‘cell leading edge’ (fig.1C). Terms relating to cell-cell adhesion were also identified (‘adherens junction’ and ‘cell-cell junction’), which may reflect the shared components between cell-cell and cell-ECM adhesion and/or the subcellular targeting of multiple baits to cell-cell contacts (e.g. as reported for vinculin, LPP, and zyxin) ^26^. Furthermore, a relatively large number of the proteins identified by proximity biotinylation were identified in published adhesomes, with 24.5% (36) and 19.7% (23) of the 147 proteins in the proximity-dependent adhesome identified in the literature-curated (232 components) and consensus adhesomes (60 components), respectively (supp. 3B) ^6, 11^. The majority of prey proteins (96; 65.3%) were also identified in at least one of the seven datasets comprising the meta adhesome. There was also substantial overlap between the prey identified for BirA*-paxillin and - kindlin-2 with those reported by Dong *et al* ^23^. (21 and 18, respectively). Together, these findings indicate that a large proportion of the proteins identified are likely to be highly relevant to IACs.

### Functional modules within IACs

Two-dimensional, hierarchical clustering was performed to provide an unbiased interrogation of the relationships between baits and the prey identified. This analysis revealed five clusters of bait proteins (B1-B5) and 16 clusters of prey (P1-P16) that are likely to represent groups of spatially-linked protein sub-complexes (fig. 2 and supp. table 2). B1 contained BirA*-tagged kindlin-2 and the members of the IPP complex (ILK, α-parvin and PINCH), B2 comprised BirA*-FAK, -paxillin, -p130Cas and -vinculin, and B3 contained BirA*-GIT1 and β-Pix. The remaining two clusters contained actin-associated/regulatory baits (BirA*-LPP and -TRIP6 in B4; BirA*-palladin, -PDLIM5, -ponsin and -zyxin in B5). These five bait clusters broadly correlated with theoretical interaction networks in the literature ^3, 11^. These findings not only provide evidence that published theoretical IAC networks are largely reflective of protein interactions *in situ*, but that BioID captures relevant interactions within IACs.

**Fig. 2.**
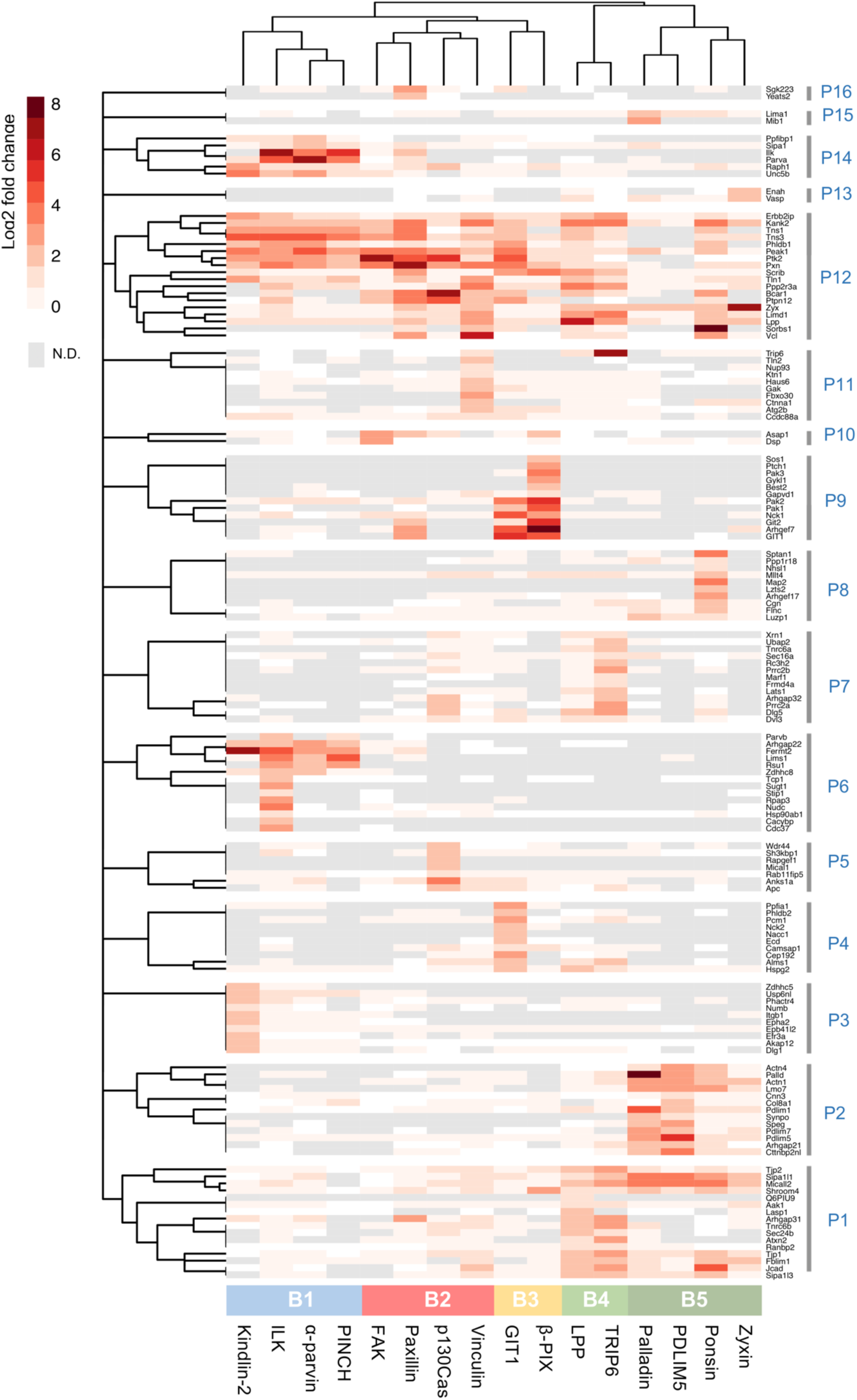
Hierarchical clustering of BioID data. Hierarchical clustering was performed on the 147 proteins identified in the proximity-dependent adhesome and displayed as a heatmap. BioID baits were clustered using the Jaccard distance of the presence (BFDR ≤ 0.05) or absence (BFDR > 0.05) of prey. Prey proteins were clustered using the Euclidean distance of log2 fold change enrichment over BirA* control. Log2 fold changes of prey proteins were displayed as a heatmap. Dendrograms were split to identify clusters of baits (B1-5, colour coded) and prey (P1-P16). N.D., not detected.

GO analysis of the prey identified by each bait revealed a number of over-represented terms relating to IACs and their associated structures. Many of these terms under the cellular component category were common to all baits, including ‘focal adhesion, ‘cell leading edge’, and ‘actin cytoskeleton’, confirming that each of the baits identified proteins relevant to cell-ECM adhesion (supp. 4A). Some GO terms were unique to a single bait, such as ‘receptor complex’ by BirA*-kindlin-2, and ‘chaperone complex’ with BirA*-ILK, suggestive of specific roles for these proteins. A broad range of GO terms was also identified across all baits under the molecular function domain, many of which were again shared (fig. 3A). For example, ‘actin binding’ was over-represented across baits in all clusters. However, other actin-related GO terms such as ‘actin filament binding’ and ‘actinin binding’ were predominantly restricted to B4 and B5, in accordance with their roles in actin regulation. Similarly, GO terms relating to GTPase binding/regulation were found by multiple baits (e.g. ‘GTPase activator activity’, ‘GTPase regulator activity’ and ‘small GTPase binding’), but were predominantly identified by B2, B3 and B4, supporting their reported roles in signalling within IACs. The similarity of GO terms identified within bait clusters indicated that these groups of proteins have similar functions, and that the clusters may have a functional relevance.

**Fig. 3.**
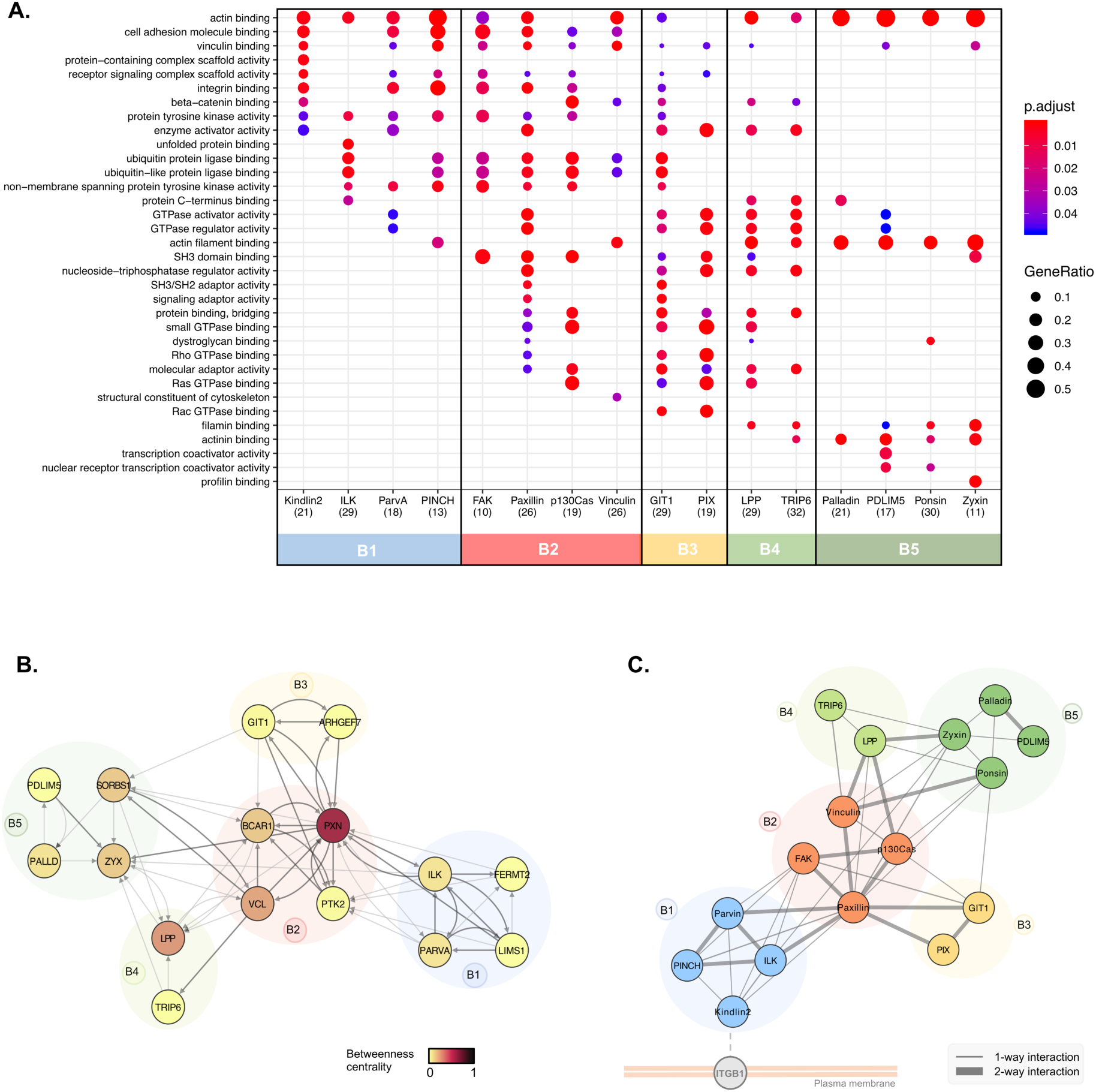
Functional roles and sub-complex organisation of functional bait modules. **A.** Functional over-representation analysis of proximal proteins identified by each of the 16 BirA*-tagged adhesome proteins (BFDR ≤ 0.05). The top 3 GO terms in the ‘molecular function’ category for each bait are listed, and displayed for all baits if identified with an adjusted p-value ≤ 0.05. The number of proteins recognised is shown in brackets. Baits are colour-coded according to hierarchical clustering shown in figure 2. p.adjust, adjusted p-value; GeneRatio, proportion of total proteins identified in each GO term. **B.** Network analysis of bait-prey interactions between BirA*-tagged baits. Nodes represent BirA*-tagged baits, which are colour-coded according to betweenness centrality, and grouped into bait clusters B1-B5, according to the hierarchical clustering (fig. 2). Edges indicate bait-prey proximity interactions, with arrow heads indicating direction of interaction (bait to prey). Dark grey edges indicate PPIs also present in a published PPI database (see methods). **C.** Schematic showing potential organisation of adhesome bait modules within IACs in relation to the membrane. Positioning of modules was guided by the functional roles of prey identified by each cluster (i.e. transmembrane protein or actin-regulatory, see supp. table 1 for details).

Close examination of the interconnectivity between baits revealed highly specific associations of bait clusters. While a large number of bi-directional proximal associations were observed within individual bait clusters, particularly for B1, B2 and B3, very few interactions were observed between clusters (e.g. no proximity interactions were observed between B1 and B3, or B1 and B4/5) (fig. 3B). The exception to this was B2, which was highly connected with components from the other clusters. Even then, connections between B2 and B1 were only mediated by paxillin and FAK, connections between B2 and B3 were via paxillin and p130Cas, and connections between B2 and B4 were via p130Cas, paxillin, and vinculin. These findings may be indicative of a spatial distinction between the clusters, and suggest that B2 may form a central link between other clusters, either in space (i.e. all modules form a single structure, with B2 in the centre) or time (i.e. B2 interacts with other bait clusters separately in different structures, such as nascent adhesions and fibrillar adhesions). Network analysis revealed that paxillin had a particularly large number of proximal interactions with BioID baits (19), and a high betweenness centrality (0.57) (fig. 3B, and supp. 2B and C) and may be indicative of a central role of paxillin as a key adaptor, mediating connections with other proteins and coordinating interactions between different bait clusters. This is consistent with the model proposed by Green and Brown, in which paxillin is described as an ’über-linker’, interacting with all functional modules ^3^. Some baits, such as PDLIM5 and β-Pix, had few connected nodes and a low betweenness centrality and are therefore likely to be more peripheral in the adhesome network. How the bait clusters identified here link to spatially-defined functions requires further investigation.

Various studies have provided evidence for functional modules within IACs, which represent groups of proteins that perform a similar function, such as signalling or mechanotransduction ^3, 11, 27^. The five bait clusters identified in this study are unlikely to represent distinct, separate structures that operate individually, and are more likely to represent dynamic, interconnected groups of proteins that interact to form IACs. Additionally, it is possible that cytosolic interactions contribute towards the network of proximity interactions described here. For example, various adhesome components have been shown to form pre-assembled dimeric or trimeric complexes in the cytosol, which are thought to facilitate the assembly of IACs in a modular manner ^28^. Indeed, many of the multimolecular interactions previously described were also identified in this study, including the trimeric complexes of ILK, PINCH and α-parvin, and FAK, p130cas and paxillin. Each of these trimers was shown to have bi-directional bait-bait proximity interactions and was present within the same bait cluster identified by hierarchical clustering.

The five bait clusters bear a striking resemblance to current models, and provide further evidence of functional modules within IACs ^3, 11^. The members of the IPP complex and kindlin-2, which form B1, are commonly grouped together in literature-based models, together with Rsu1 (a prey identified by BirA*-tagged ILK, PINCH and α-parvin in this study). Although the exact role of this functional module is not fully understood, it has been proposed to play a role in the recruitment of regulatory components to IACs to regulate Rac1 GTPase activity and other signals at the cell cortex ^3, 29^. B2 comprises well-studied IAC regulators and adaptors (FAK, paxillin, vinculin, and p130Cas). Each of the baits within this highly connected central cluster has been shown to play a role in mechanotransduction, and future experiments examining the effects of force on the proximity-dependent adhesome will be instructive ^30–32^. The components of B3, GIT1 and β-Pix, have been shown to form a stable complex, and regulate various signalling pathways and cytoskeletal dynamics via effectors such as Rho family GTPases and PAK family kinases (prey identified by BirA*-GIT1 and -β-Pix in this study) ^33^. Although they are often treated as a single entity, GIT and Pix have different interacting partners and there is evidence that they function independently of one another ^33^. This may account for the differential proximal interactors identified by each component in this study. The final two bait clusters represent two actin-regulatory modules, and contain a number of components responsible for mediating the connection between actin filaments and IACs through the recruitment of actin regulatory and bundling proteins, such as α-actinin. The inclusion of additional BioID baits, such as integrins, talin and tensins, may provide useful information about the interconnectivity between these modules through the detection of connecting proteins.

### Substructure and stratification of the proximity-dependent adhesome

Although super-resolution microscopy has revealed the organisation of a small number of components within IACs, little is known about the localisation of the majority of IAC components, and a more comprehensive view could provide insights into IAC regulation and signalling outputs. A key advantage of BioID is that it provides a means to interrogate spatial relationships between groups of proteins. Examination of the prey identified by each bait cluster may therefore provide insights into the spatial organisation of IACs, and build upon previous evidence for sub-structure ^14, 23^. As multiple membrane-associated proteins and transmembrane receptors, such as integrin β1 (Itgb1) and netrin receptor (Unc5b), were predominantly identified by BirA*-kindlin-2 and other members of B1 (supp. table 1), it is likely that these baits lie within close proximity to the plasma membrane (fig. 3C). Conversely, the large proportion of actin-regulatory proteins detected by the actin-associated baits in B4 and B5 indicates that these proteins lie more distal from the membrane, in the proximity of actomyosin filaments. As members of B2 have multiple links to B1 and B4/B5, these proteins may lie within or between these two regions. This organisation broadly correlates with the stratified architecture of IACs determined by super-resolution microscopy ^14^, with zyxin (B5) and prey identified by B4 and B5 (VASP and α-actinin) localised to a membrane-distal actin-regulatory layer, and members of B2 (FAK, paxillin, and vinculin) distributed across the force-transduction layer and integrin-signalling layer. Although the stratified organisation of members of B1 has not yet been examined in mammalian cells in culture, recent work in Drosophila has localised them to the membrane-proximal integrin signalling layer ^34^.

The stratified organisation of components determined by BioID is further supported by the detection of biotinylation within specific domains of talin by each of the BioID baits (supp. table 3). Talin has a polarised orientation within IACs and spans the three layers of the stratified model of IAC architecture, with its N-terminal FERM domain being located proximal to the plasma membrane, and the end of its C-terminal rod domain mediating attachment to actin filaments ^14, 18^. The peptides within talin that were biotinylated by its associating baits were mapped to its primary and tertiary structure (supp. 3A and B), and found to correlate with the organisation of bait clusters relative to the plasma membrane outlined in fig. 3C. The actin-associated baits BirA*-LPP, -TRIP6, -zyxin and -ponsin, and BirA*-vinculin, biotinylated peptides in the C-terminal R11 and R13/DD domains of talin, which lie proximal to the ABS3 actin-binding site ^35^. Despite the many reported vinculin binding sites in talin, and localisation to multiple layers of IACs^15^, vinculin only biotinylated peptides at the C-terminus of talin ^35^. No biotinylated peptides from B4 or B5 baits were found in the ABS1 or ABS2 domains of talin, despite their reported roles in actin binding ^36–38^. By contrast, B1 baits biotinylated peptides located in the membrane-proximal linker domain of talin, close to the IBS1 integrin-binding site and consistent with direct kindlin-integrin binding ^39^. BirA*-paxillin biotinylated peptides from various domains across the length of talin, which may indicate that paxillin is localised across multiple layers of IACs. Alternatively, these may represent interactions with talin in its autoinhibited, inactive conformation ^40^. Paxillin has been reported to interact with the talin R7/R8 domain via its leucine-aspartic acid (LD) domains ^41^, though it is likely that there are additional binding sites (B. Goult, personal communication, January 2020).

Evidently, inferring IAC sub-structure from proximity-dependent labelling relies on making a number of assumptions (i.e. shared proximal interactions occur in the same time and place), and further experiments are required to confirm such speculations. Nevertheless, despite capturing interactions from a heterogenous population of IACs, the organisation of adhesome proteins inferred from proximity biotinylation correlates with models of IAC architecture, such as the stratified organisation of IACs in mammalian cells and myotendinous junctions in *Drosophila* ^14, 34^. The specific biotinylation of talin domains by BioID baits provides further evidence for a high degree of organisation. Although the stratified arrangement of BioID baits needs to be further validated, this data could be used to infer the localisation of prey proteins based on the baits with which it was proximally associated.

### Topological organisation of the proximity-dependent adhesome

The hierarchical cluster analysis of bait proteins and prey was then used to interrogate the topology of the proximity-dependent adhesome network, and GO analysis performed to determine the functional relevance of prey clusters (fig. 4 and supp. 2, 4A and B). Organisation of the network was driven by the hierarchical cluster analysis, with baits excluded from the prey clusters. While some proteins had shared interactions with multiple baits and bait clusters, others were uniquely identified by a single bait, and indicate underappreciated links to more distal roles. For example, a subgroup of eight proteins in prey cluster P6, exclusively identified by BirA*-ILK, contains a large number of Hsp90-binding chaperone/co-chaperone proteins. The association of ILK with Hsp90 has been previously reported ^42^, but this link may be more significant for IACs than previously thought. Similarly, BirA*-GIT1 uniquely identified a number of microtubule-associated proteins, with over-represented GO terms such as ‘microtubule binding’ in the molecular function category and those relating to the centriole in the cellular component category. These associations are in line with the reported role of GIT1 in microtubule nucleation at the centrosome ^43, 44^. Furthermore, BirA*-TRIP6 has a number of unique links to proteins involved in RNA binding and regulation, and the over-represented GO terms from P7 under the cellular component category include ‘P-body’. TRIP6 has been reported to localise to the nucleus, and other zyxin family members LIMD1, ajuba and WTIP were shown to associate with processing-bodies (P-bodies) in U2OS cells ^45^. Although TRIP6 itself was shown to have poor colocalisation with P-bodies, it is possible that zyxin plays a role in RNA regulation.

**Fig. 4.**
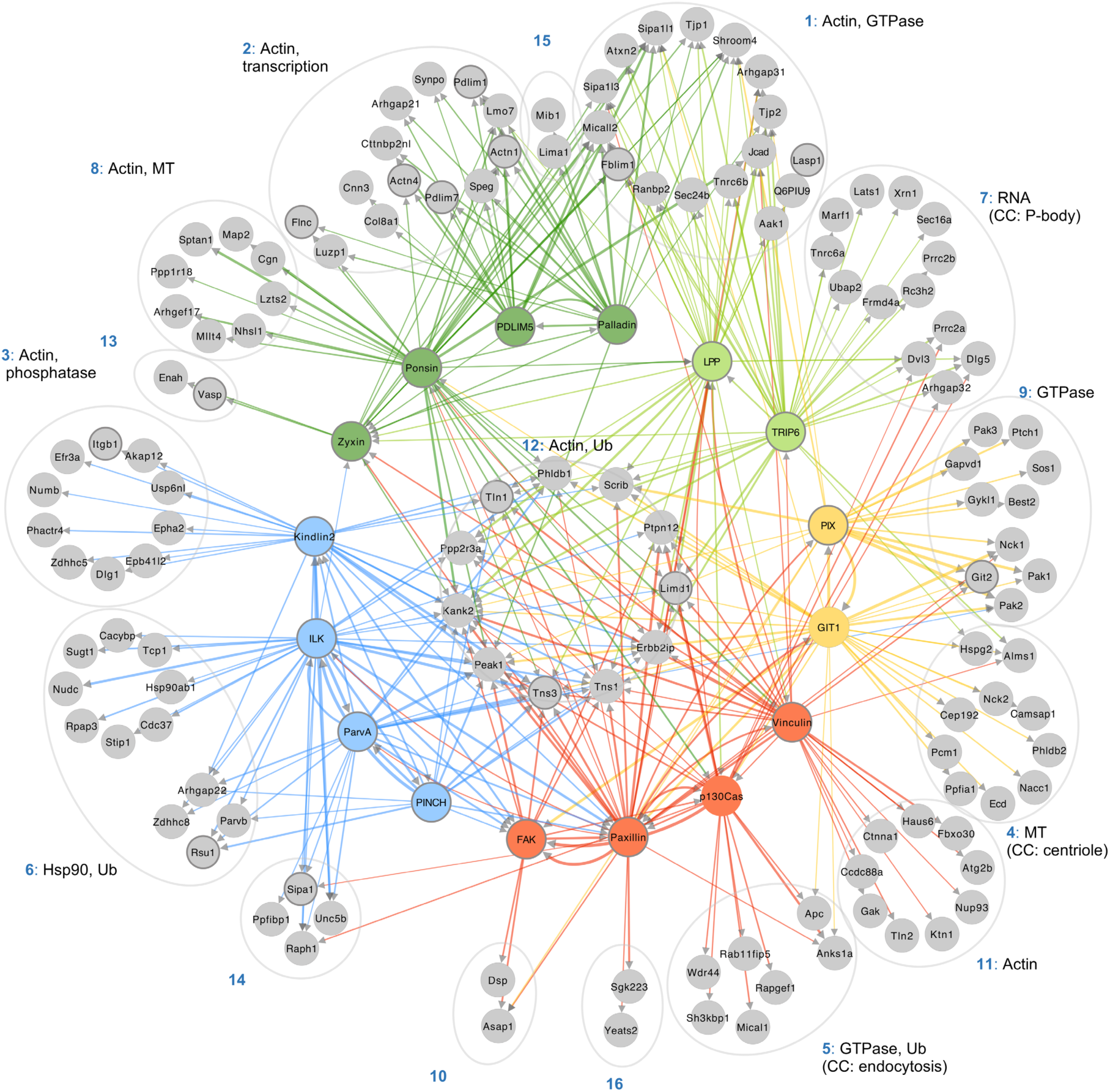
Topological organisation of the proximity-dependent adhesome. Network of proximity interactions within the proximity-dependent adhesome. Network organisation was driven by hierarchical clustering of BirA*-tagged adhesome baits and proximal prey proteins (BFDR ≤ 0.05) (fig. 2). Grey nodes represent prey proteins, and nodes indicating BirA*-tagged adhesome bait proteins are colour-coded according to the hierarchical clustering in figure 2. Consensus adhesome components are indicated with thick grey outlines11. Edges indicate bait-prey proximity interactions, with colour representing source node and width representing fold-change over BirA* control. The top GO terms under the molecular function category for each prey cluster are indicated. Gene names are shown. CC, cellular component; MT, microtubule; Ub, ubiquitination.

Other prey clusters were associated with multiple bait proteins. The actin-associated baits in B4 and B5 were highly connected to prey in P1 and P2. GO terms relating to actin regulation were well represented within these clusters, and may be indicative of an actin filament regulatory module. Furthermore, a central group of 11 highly-connected proteins (P12) had multiple links to all five bait clusters, and may represent core IAC components. Multiple well-established adhesome proteins were identified within this central group, including talin-1 and tensins-1 and −3, in addition to the more recently identified IAC component, KANK2, and the cortical microtubule stabilising complex (CMSC) component LL5-α (Phldb1) ^46, 47^.

KANK2 was robustly identified by almost all the BioID baits in this study, and as a proximity interactor of both paxillin and kindlin-2 by proximity biotinylation in U2OS cells ^23^. The role of KANK proteins in cell-ECM adhesion has become apparent in recent years, as they have been shown to be involved in the turnover of IACs through the recruitment of CMSCs to IACs and the uncoupling of mechanical transduction between integrins and the actomyosin network, resulting in sliding focal adhesions ^47, 48^. Although KANK is known to bind talin, a direct interaction with any of the baits used in this study has yet to be described ^47^. As KANK2 was identified in a number of proximity interactomes, it is possible that additional direct interactions exist^19^. In addition to KANK2 and LL5-α (Phldb1), three other CMSC components were identified in the proximity-dependent adhesome, LL5-β (Phldb2), liprin-α1 (Ppfia1) and liprin-β1 (Ppfibp1). This provides further evidence of the association of CMSCs and microtubule-associated structures with IACs ^47, 49, 50^. Some of these components exhibited a restricted set of binding partners. For example, liprin-α1 and LL5-β were uniquely linked to BirA*-GIT1, and liprin-β1 was uniquely detected by BirA*-α-parvin, which may indicate specific roles for GIT1 and α-parvin in CMSC regulation and microtubule-targeting to IACs.

Peak1, PTP-PEST (Ptpn12), and LIMD1 have previously been identified as adhesome components, though their precise roles in IAC regulation are less well studied ^6, 11, 51–53^. Peak1 (also known as SgK269), is a pseudokinase that functions as a scaffolding protein to recruit various signalling molecules, and its overexpression has been linked to progression of various cancers, including pancreatic ductal adenocarcinoma ^54–56^. Peak1 localises to IACs and the actin cytoskeleton following growth factor receptor stimulation and cellular attachment, where it regulates cell motility, spreading, and IAC turnover ^51, 57^. The signalling activity of Peak1 is mediated by phosphorylation by Src family kinases, such as Src and Lyn ^54, 55^. In turn, Peak1 regulates phosphorylation of adhesome components such as paxillin and p130Cas, though the mechanisms are unknown ^51, 57^. Indeed, both BirA*-paxillin and -p130Cas detected Peak1 as a proximal interactor. Peak1 is also identified by many other baits, and may act as an adaptor to recruit other adhesome components.

Other proteins within the central cluster have very few reported associations with IACs, including scribble (Scrib), erbin (Erbb2ip), and the PP2A phosphatase regulatory subunit, Ppp2r3a. These proteins may represent underappreciated IAC components and regulators. Scribble is known as an adaptor protein that regulates cell polarity, but it has also been reported to interact with a number of adhesome components, including LPP, TRIP6 and β-Pix, and co-immunoprecipitate with others, such as GIT1, Pak and integrin α5 ^58–62^. Consistent with these reports, scribble was identified as a proximal interactor by BirA*-tagged LPP, TRIP6, β-Pix and GIT1 (together with kindlin-2 and paxillin). Although scribble is typically localised to cell-cell junctions, a number of studies have reported its recruitment to the leading edge of migrating non-fibroblastic cells, where it colocalised with β-Pix and Cdc42 and regulated directional cell migration ^58, 59, 63, 64^. It is therefore conceivable that scribble also plays a role in IACs and directional cell migration in fibroblasts. Scribble was also detected in the meta adhesome (2 of 7 datasets), and was identified in a phosphoproteomic analysis of IACs ^11, 65^. Similar to scribble, erbin also localises to the basolateral membrane at cell-cell junctions in epithelial cells, and is also found at synapses ^66, 67^. Other than evidence for an interaction of erbin with integrin β4 in hemidesmosomes, it has few published links to adhesome components, and a potential role in the IAC regulation has yet to be explored ^68^. Like scribble, erbin has been localised to the leading edge of cells, and was identified in the meta adhesome (1 of 7 datasets) ^11, 69, 70^. However, considering erbin is identified as a proximal interactor by a different subset of adhesome baits, and is not detected by BirA*-β-Pix and -GIT1, it is likely that the two proteins have different binding partners and perform different roles.

Two phosphatases, PTP-PEST and a regulatory subunit of PP2A (Ppp2r3a), were also identified, and may represent underappreciated core regulators of IAC dynamics and signalling outputs. PTP-PEST, a tyrosine phosphatase, is reported to have a key role in IAC turnover and cell motility through the dephosphorylation of a number of core adhesome components, including p130Cas, FAK, and paxillin ^71–73^. In line with this, PTP-PEST was identified as a proximal interactor by BirA*-tagged paxillin, FAK, and p130Cas. PTP-PEST was also identified as a proximal interactor by BirA*-vinculin, -ILK and -GIT1, though these proteins have not been identified as substrates for PTP-PEST, and may represent indirect interactors. Ppp2r3a, a regulatory subunit of the serine threonine phosphatase heterotrimer PP2A, is part of the PR72/PR130 subgroup of PP2A isoforms ^74^. Recently, ppp2r3a was shown to regulate cell migration via interaction with LPP LIM domains ^75^. Although it colocalised with LPP at the cell periphery in spreading cells, ppp2r3a was excluded from mature IACs. It is thought that LPP may bind ppp2r3a to target PP2A to early IACs, bringing it within close proximity to enable dephosphorylation of substrates to regulate dynamic IAC turnover and enable effective cell migration. However, although it is well-established that PP2A can regulate IACs via dephosphorylation of paxillin ^76^, the potential PR72/PR130 family-specific PP2A substrates have yet to be identified. In this study, ppp2r3a was identified as a proximal interactor by BirA*-tagged LPP, TRIP6, vinculin, p130Cas, GIT1, and ILK. Whether these represent PP2A substrates or adaptor proteins that recruit PP2A to IACs via ppp2r3a is unknown. Recently, ppp2r3a and LPP were identified in a proteome-wide screen to identify novel LD motifs ^77^. The LD motifs in paxillin interact with various adhesome proteins containing LD binding domains, including ILK, vinculin, GIT, and talin, among others, and it is feasible that the LD motifs in ppp2r3a and LPP also facilitate interactions with such proteins ^41, 78–80^.

A closer examination of the relative abundances of the central components revealed differential detection. While some proteins were detected at relatively similar levels across baits in a range of clusters (e.g. KANK2, Peak1 and talin), others were detected with high relative abundance by one or two baits within a single cluster (fig. 5). For example, PTP-PEST was detected with high relative abundance by BirA*-p130Cas and -paxillin, which may suggest a more specific association. Both paxillin ^81^ and p130Cas ^82, 83^ have been reported to be substrates for PTP-PEST. Some prey proteins showed similar patterns of bait ID and relative abundance (e.g. Ppp2r3a/LIMD1, and Kank2/Peak1/Tns1/Tns3), which may suggest that these proteins have similar roles. Future experiments examining the role of these central components in IAC function and regulation will be informative.

**Fig. 5.**
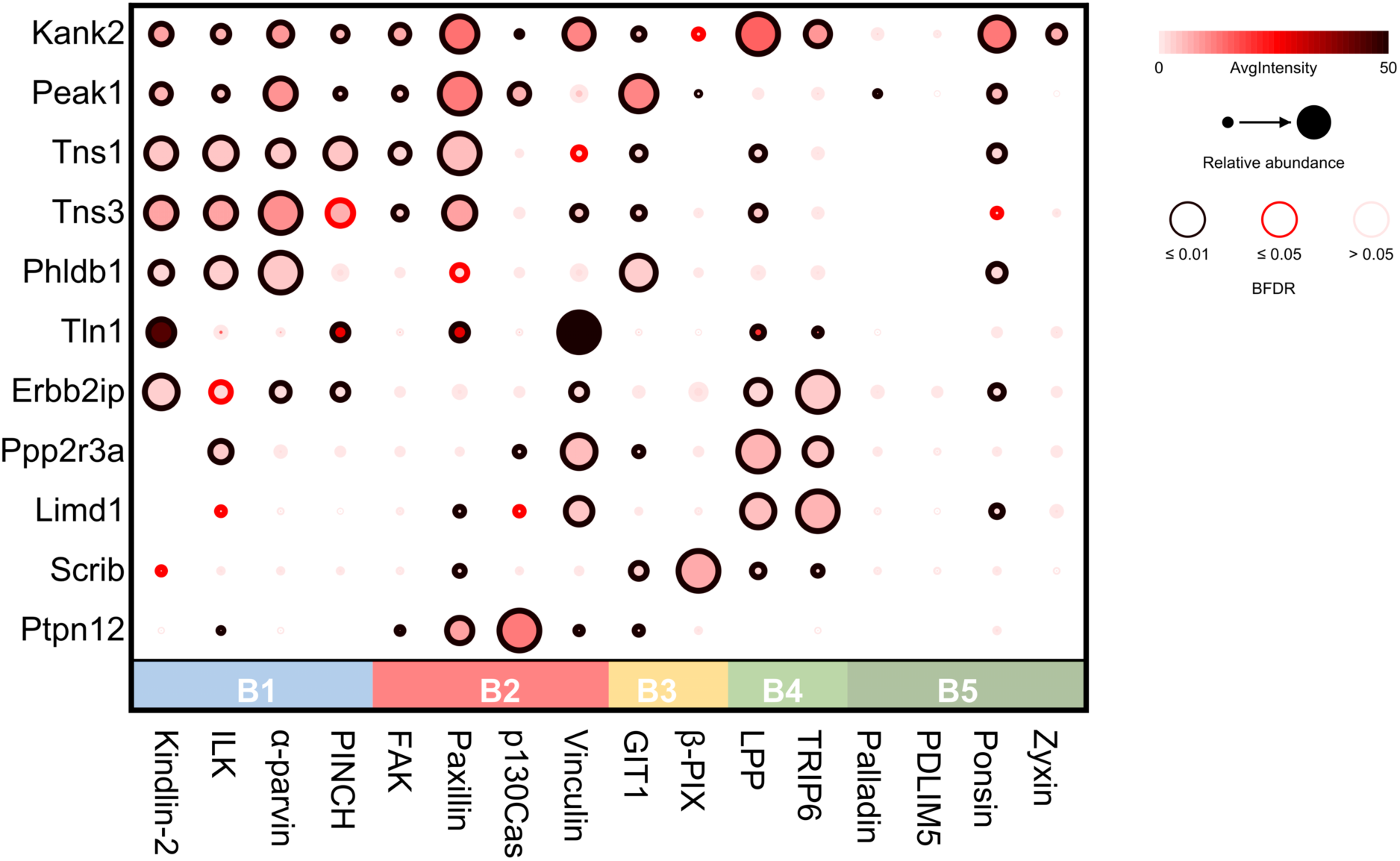
Proximity interactions of the central prey cluster. Dot plot of proteins within the central prey cluster (P12) of the proximity-dependent adhesome. Prey also used as baits found in the central cluster were excluded (Ptk, Pxn, Vcl, Bcar1, Zyx, Lpp, Sorbs1). Baits were organised into clusters defined by hierarchical clustering in figure 2. AvgIntensity, average intensity (generated from SAINTexpress); BFDR, Bayesian false discovery rate. Dot plot was generated by Prohits-Viz 84.

## Summary

In this study, multiplexed proximity-dependent biotinylation was used to generate an empirically-defined network of proximal associations within IACs. Unbiased bioinformatic analysis of the network revealed five groups of baits which link to common groups of prey, and may represent functional modules within IACs. The interconnectivity between these modules and their inferred stratified organisation are consistent with current models of adhesome PPI networks and IAC architecture. A large number of IAC-associated proteins were identified within the proximity-dependent adhesome, in addition to a range of prey that may represent novel IAC components, or underappreciated links to other cellular organelles. Due to the multi-functionality of many of the BioID baits, it is not possible to conclude that proximal interactors associate with IACs directly, or localise to more distal structures without further experiments. For example, vinculin, LPP, TRIP6 and zyxin also localise to cell-cell contacts, and it is possible that a proportion of the proximity interactions identified lie at cell-cell junctions. Nonetheless, a central group of 11 highly-connected prey was identified which may represent core adhesome components, some of which have few reported associations with IACs. The detection of these proteins by a number of adhesome baits suggests that they may play a more central role in IACs than currently appreciated, and future studies should focus on their role.

While proximity-dependent labelling methods, such as BioID, have become increasingly popular to examine individual protein interactomes and in large-scale initiatives to map protein interaction networks, there are limitations that must be kept in mind when interpreting data. For example, although highly stringent analyses were performed, it is possible that a number of non-specific contaminants were identified. For example, despite being extracellular proteins, perlecan (Hspg2) and collagen alpha-1 (VIII) chain (Col8a1) were identified in the proximity-dependent adhesome and are likely to represent false positives. Similarly, some proximal interactors may be missed due to the restricted labelling radius of BirA* (10-15 nm) and the dependency of labelling on accessible lysines. Indeed, some reported adhesome components were not identified in this study, such as Csk and Hic-5. Finally, due to the differential availability of lysine residues, protein turnover rates and mass spectrometric detection of individual peptides and proteins, proximity-dependent biotinylation is unable to differentiate between the degree of proximity, protein abundance, or frequency/permanence of interactions.

Nevertheless, this study has generated insights into the topological organisation of the adhesome and has highlighted some underappreciated components that may play a central role in IAC function and regulation. The study therefore provides a useful resource to drive further hypothesis generation, and demonstrates that proximity-dependent labelling is a valuable addition to the tools currently available to examine IAC composition and protein-protein relationships. Future studies that focus on how this network is altered under disease-relevant conditions (e.g. under different force conditions ^85^ or throughout the cell cycle ^86, 87^) may further our understanding of the role of IACs in governing cellular behaviour in health and disease.

## Methods

### Reagents

All reagents were acquired from Sigma-Aldrich (St. Louis, MO) unless otherwise specified. Primary antibodies used for immunofluorescence microscopy were mouse anti-vinculin (hVin-1, Sigma; 1:200), rabbit anti-paxillin (GeneTex, Irvine, CA; 1:200), mouse anti-c-myc (9B11, Cell Signalling Technologies, Danvers, MA; 1:200). Alexa Flour 680-conjugated streptavidin was from Life Technologies, and secondary antibodies (anti-mouse IgG Alexa Flour 488 and anti-rabbit IgG Alexa Flour 488) were from Invitrogen (Carlsbad, CA).

### Cell culture

Mouse pancreatic fibroblasts (im-PSC ^88^) and HEK 293 cells were cultured in D5796 Dulbecco’s-modified Eagle’s medium (DMEM) supplemented with 10% (v/v) fetal bovine serum (FBS; Life Technologies, Carlsbad, CA) and 2 mM L-glutamine. Cells were maintained at 37°C in a humidified atmosphere with 5% (v/v) CO_2_.

### Cloning

The BioID vectors pCDH-TagBFP-T2A-myc-BirA* ^89^ and pCDNA3.1-BirA*-paxillin were gifts from A. Gilmore (University of Manchester) and E. Manser (IMCB, A*STAR, Singapore). The plasmids containing ILK, ponsin, kindlin-2, vinculin ^90^ and α-parvin were a gift from C. Ballestrem (University of Manchester) and the plasmid containing zyxin was a gift from A. Sharrocks (University of Manchester). The pcDNA3.1-myc-BirA*-LPP and pcDNA3.1-myc-BirA*-TRIP6 plasmids from which LPP and TRIP6 were amplified, respectively, were generated by J. Askari and J. Zha (University of Manchester) from plasmids containing LPP and TRIP6 that were a gift from A. Sharrocks (University of Manchester). Flag-ECFP-betaPixa (plasmid #15235), mEmerald-PINCH-C-14 (plasmid #54229), mCherry-Palladin-C-7 (plasmid #55113), pEGFP-GIT1 (plasmid #15226) and pGFP-Cas (plasmid #50729) were purchased from Addgene. Full length open reading frames (ORFs) of target adhesome proteins were amplified by polymerase chain reaction, and cloned into the pCDH-TagBFP-T2A-myc-BirA* vector using Gibson assembly (vinculin, ponsin and p130Cas), HiFi DNA assembly (FAK, kindlin-2, β-Pix, palladin, α-parvin, PINCH, PDLIM5 and zyxin), or restriction enzymes (BirA*-paxillin, BirA*-LPP, BirA*-TRIP6; BspEI and SalI-HF, BirA*-ILK; XhoI and SalI-HF (see supp. table 4 for primer pairs and annealing temps). During PCR amplification, two different annealing temperatures were used to promote efficient primer annealing first to the plasmid template (10 cycles) then PCR product template (25 cycles). All constructs included a five amino acid linker (LERPL) between BirA* and the protein of interest. Primers for Gibson assembly and HiFi assembly were designed using SnapGene (GSL Biotech LLC, Chicago, IL), and primers were manufactured by Integrated DNA Technologies (Newark, NJ). ORF sequences were confirmed by sequencing.

### Generation of stable cell lines

Lentiviruses containing BirA* constructs were produced by transient co-transfection of HEK 293 cells with pCDH-TagBFP-T2A-myc-BirA* plasmids and packaging vectors (psPAX2 and pM2G) using polyethylenimine (PEI)-mediated transfection. 250 μl DNA mix containing 6 μg pCDH-TagBFP-T2A-myc-BirA* plasmid, 4.5 μg psPAX2 and 3 μg pM2G in Opti-MEM reduced serum media (Thermo Fisher, Waltham, MA) was added to 250 μl PEI mix (44.4 μM PEI, 1.5 mM NaCl in Opti-MEM) and incubated at room temperature (RT) for 20 minutes. HEK 293 cells (T75 flask, ∼60% confluency) were incubated with 5 ml Opti-MEM and PEI/DNA mix for 6 hours before medium replaced with fresh medium. Three days post-transfection, filter-sterilised viral medium was added to im-PSC cells for 24 hours before being replaced with fresh medium, and cells passaged 24 to 48 hours later. Cells expressing blue fluorescent protein were selected using fluorescence activated cell sorting, and sorted into high, medium, and low-expressing populations. Western blotting and immunofluorescence microscopy were used to confirm expression of full-length constructs and select appropriate cell populations with clear subcellular targeting of bait proteins (and biotinylated proteins) to IACs with minimal background localisation and biotinylation for use in subsequent experiments.

### Proximity biotinylation and affinity purification of biotinylated proteins

To induce proximity biotinylation, cells expressing BirA* constructs were seeded onto plastic tissue culture plates for 8 hours to allow for robust IAC formation, then incubated in medium with 50 μM biotin for 24 hours. Biotinylated proteins were affinity purified following a protocol adapted from Roux *et al*., 2016 ^91, 92^. Three 10 cm plates of cells were washed three times in PBS- and cells lysed with 400 μl lysis buffer (50 mM Tris/HCl, pH 7.4, 250 mM NaCl, 0.1% (w/v) SDS, 0.5 mM DTT, 1X cOmplete^TM^ Protease Inhibitor Cocktail) at RT. 120 μl 20% (v/v) Triton X-100 was added, and cell lysates maintained at 4°C. DNA was sheared by passing cell lysates through a 19 G needle four times and through 21 G needle four times before 360 μl chilled 50 mM Tris-HCl pH 7.4 added, and passed through a 27 G needle four times. Cell lysates were centrifuged at full speed for 10 minutes at 4°C, and supernatant rotated with 45 μl MagReSyn^®^ streptavidin beads (ReSyn Biosciences, Gauteng, South Africa) at 4°C overnight. Beads were washed twice with 500 μl wash buffer 1 (10% (w/v) SDS), once with 500 μl wash buffer 2 (0.1% (w/v) deoxycholic acid, 1% (w/v) Triton X-100, 1 mM EDTA, 500 mM NaCl, 50 mM HEPES), and once with 500 μl wash buffer 3 (0.5% (w/v) deoxycholic acid, 0.5% (w/v) NP-40, 1 mM EDTA, 10 mM Tris/HCl, pH 7.4). Proteins were eluted in 100 μl 2X reducing sample buffer with 100 μM biotin for 10 minutes at 70°C. The presence of biotinylated proteins was confirmed using western blotting, and samples analysed using liquid chromatography tandem MS (LC-MS/MS).

### Mass spectrometry sample preparation

Eluted proteins were briefly subjected to SDS-PAGE (3 minutes at 200 V, 4-12% Bis-Tris gel, Life Technologies), and stained with InstantBlue^TM^ Coomassie protein stain before being washed with ddH2O overnight at 4°C. Bands were excised and transferred to wells in a perforated 96-well plate, and in-gel tryptic digestion performed, as previously described^12^. Peptides were desalted using 1 mg POROS Oligo R3 beads (Thermo Fisher). Beads were washed with 50 μl 0.1% (v/v) formic acid (FA) before the peptide solution was added. Beads were washed twice with 100 μl 0.1% (v/v) FA, and peptides eluted with 50 μl 50% (v/v) acetonitrile (ACN), 0.1% (v/v) FA, twice. Peptides were dried using a vacuum centrifuge and resuspended in 11 μl 5% (v/v) ACN, 0.1% (v/v) FA, before analysis by LC-MS/MS.

### Mass spec data acquisition

Peptides were analysed using LC-MS/MS using a 3000 Rapid Separation LC (RSLC, Dionex Corporation, Sunnyvale, CA) Q Exactive^TM^ HF mass spectrometer (Thermo Fisher). Mobile phase A was 0.1% (v/v) FA in water, and mobile phase B was 0.1% (v/v) FA in ACN, and a 75 mm x 250 μm i.d. 1.7 mM CSH C18 analytical column (Waters, Milford, MA) was used. 3 μl of sample was transferred to a 5 μl loop and loaded on to the column at a flow rate of 300 nl/min for 13 minutes at 5% (v/v) mobile phase B. The loop was taken out of line and the flow reduced to 200 nl/min in 30 seconds. Peptides were separated using a gradient of 5% to 18% B in 34.5 minutes, then from 18% to 27% B in 8 minutes and 27% to 60% B in 1 minute. The column was washed at 60% B for 3 minutes before re-equilibration to 5% B in 1 minute. Flow was increased at 55 minutes to 300 nl/min until the end of the run at 60 min. Mass spectrometry data were acquired in a data-directed manner for 60 min in positive mode. Peptides were automatically selected for fragmentation by data-dependent analysis on a basis of the top 12 peptides with m/z between 300 to 1750Th and a charge state of 2, 3, or 4 with a dynamic exclusion set at 15 sec. The MS resolution was set at 120,000 with an AGC target of 3e6 and a maximum fill time set at 20 ms. The MS2 resolution was set to 30,000 with an AGC target of 2e5, a maximum fill time of 45 ms, isolation window of 1.3Th and a collision energy of 28.

Raw data were processed using MaxQuant (v1.6.2.10, available from Max Planck Institute of Biochemistry) ^93^. All experiments using mouse BioID baits were searched against the mouse proteome obtained from Uniprot (August 2018) ^94^. Experiments involving non-mouse BioID baits were run individually against the same mouse proteome with the relevant non-mouse BioID bait protein sequence appended. Default parameters were used in MaxQuant, with the addition of biotinylation of lysine as a variable modification, match between runs turned on, LFQ quantification selected and unique peptides only for protein quantification. All mass proteomic data are available via ProteomeXchange with identifier PXD017241.

### Bioinformatic analyses

MaxQuant protein LFQ intensities were used to assess the confidence of bait-prey interactions by MS1 intensity-based SAINTexpress ^95^ (v3.6.3). Default parameters were used, and a BFDR of ≤ 0.05 used as a stringent threshold to identify high confidence bait-prey proximity interactions.

Pairwise comparisons, hierarchical cluster analyses, and visualisation of talin biotinylated peptides were performed in R. Hierarchical clustering of baits was performed using the Jaccard distance of proximal prey (BFDR ≤ 0.05, present; BFDR > 0.05, absent), and prey were clustered using the Euclidean distance of fold-change enrichment over control. Results were displayed as a hierarchically clustered heatmap (log_2_ fold-change values visualised).

Network visualisation and analyses were performed using Cytoscape (v3.7.1) ^96^. Proteins were mapped onto an interaction network compiled from mouse, rat and human interaction databases from the Biological General Repository for Interaction Datasets (BioGRID; 3.5.166, November 2018), the MatrixDB (April 2012), and the literature-curated adhesome ^6, 97, 98^. Network analysis was performed using the NetworkAnalyzer plugin in Cytoscape ^99^. GO analyses were performed and visualised using the clusterProfiler package in R ^100^. Biotinylated peptides were searched against the mouse talin-1 sequence from UniProt (P26039) to identify biotinylated lysine sequence positions. Highly confident biotinylated lysine positions were selected for mouse talin-1 with MaxQuant localisation probability > 0.75. The dot plot in fig. 5 was generated using ProHits-Viz^84^, using the average intensity generated by SAINTexpress as a measure of abundance.

### Immunofluorescence microscopy

Cells expressing BirA* constructs were plated onto glass coverslips for 24 hours and incubated with 50 μM biotin for 24 hours to initiate biotinylation of proximal proteins. Cells were fixed with 4% (w/v) paraformaldehyde for 15 minutes at RT, and permeabilised with 300 μl 0.2% (w/v) Triton X-100 for 20 minutes at RT. Coverslips were incubated with primary antibodies directed against proteins indicated in 2% (w/v) bovine serum albumin (BSA) in PBS-for 1 hour at RT. Cells were then incubated with fluorophore-conjugated secondary antibodies at RT for 20 minutes, and stained with 1 μg/ml DAPI for 1 minute before washing and mounting onto glass slides. Images were acquired using an Olympus BX51 upright microscope with a 60x/0.65-1.25 UPlanFLN or 10x/0.30 UPlanFLN objective and captured using a Coolsnap EZ camera (Photometrics, Tucson, AZ) through MetaVue software (Molecular Devices, San Jose, CA).

### Data deposition

The mass spectrometry proteomics data have been deposited to the ProteomeXchange Consortium via the PRIDE partner repository with the dataset identifier PXD017241^101^.

## Acknowledgments

This work was supported by a Cancer Research UK Programme Grant (C13329/A21671; to MJH), Institute Award (A19258; CJ) and Experimental Medicine Programme Award (A25236; to CJ), and by the Rosetrees Trust (M286; to CJ) and a European Research Council Consolidator Award (ERC-2017-COG 772577; to CJ). Megan Chastney was supported by a PhD studentship from BBSRC.

## Supplementary figures

**Supp. 1.**
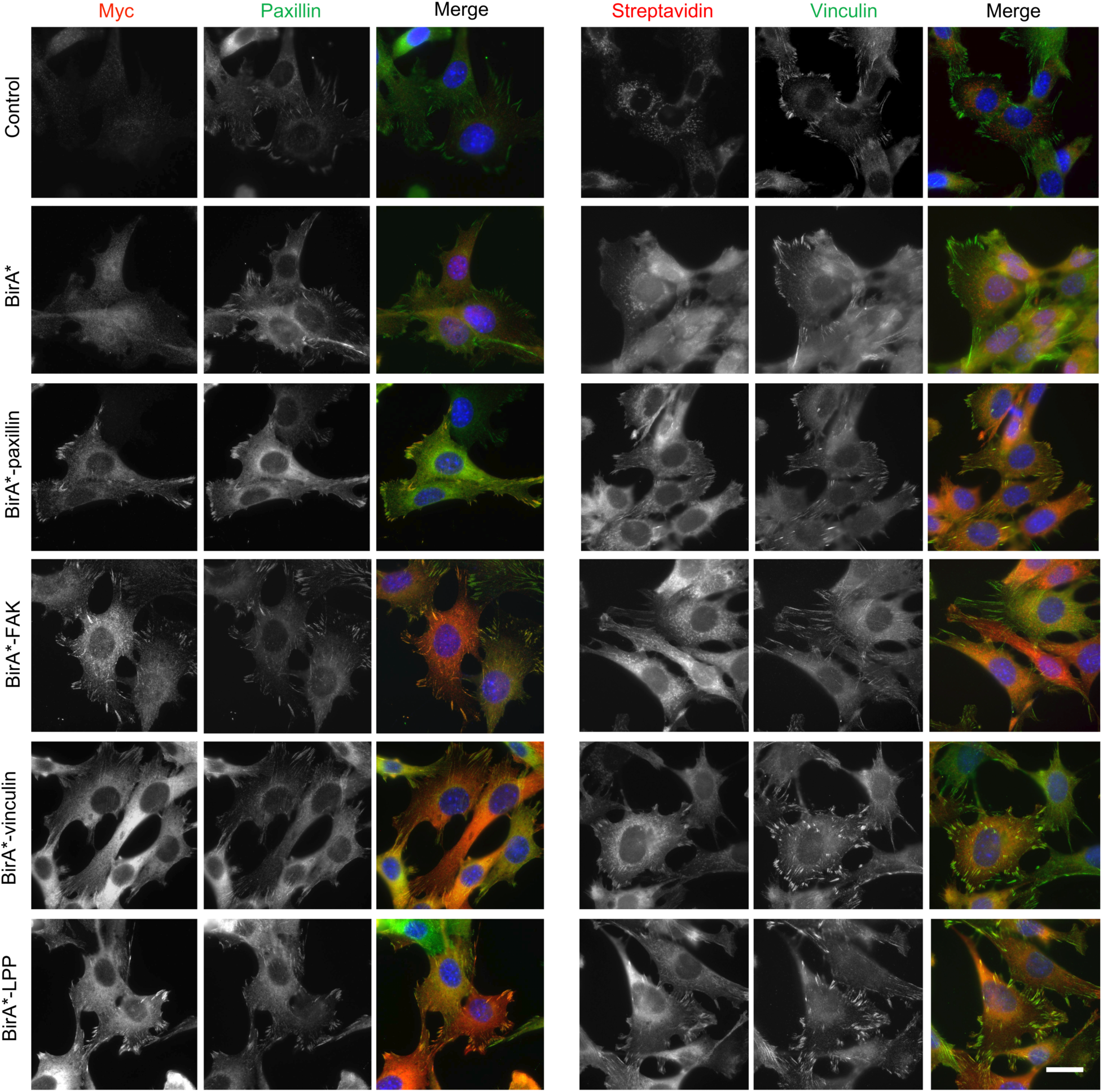
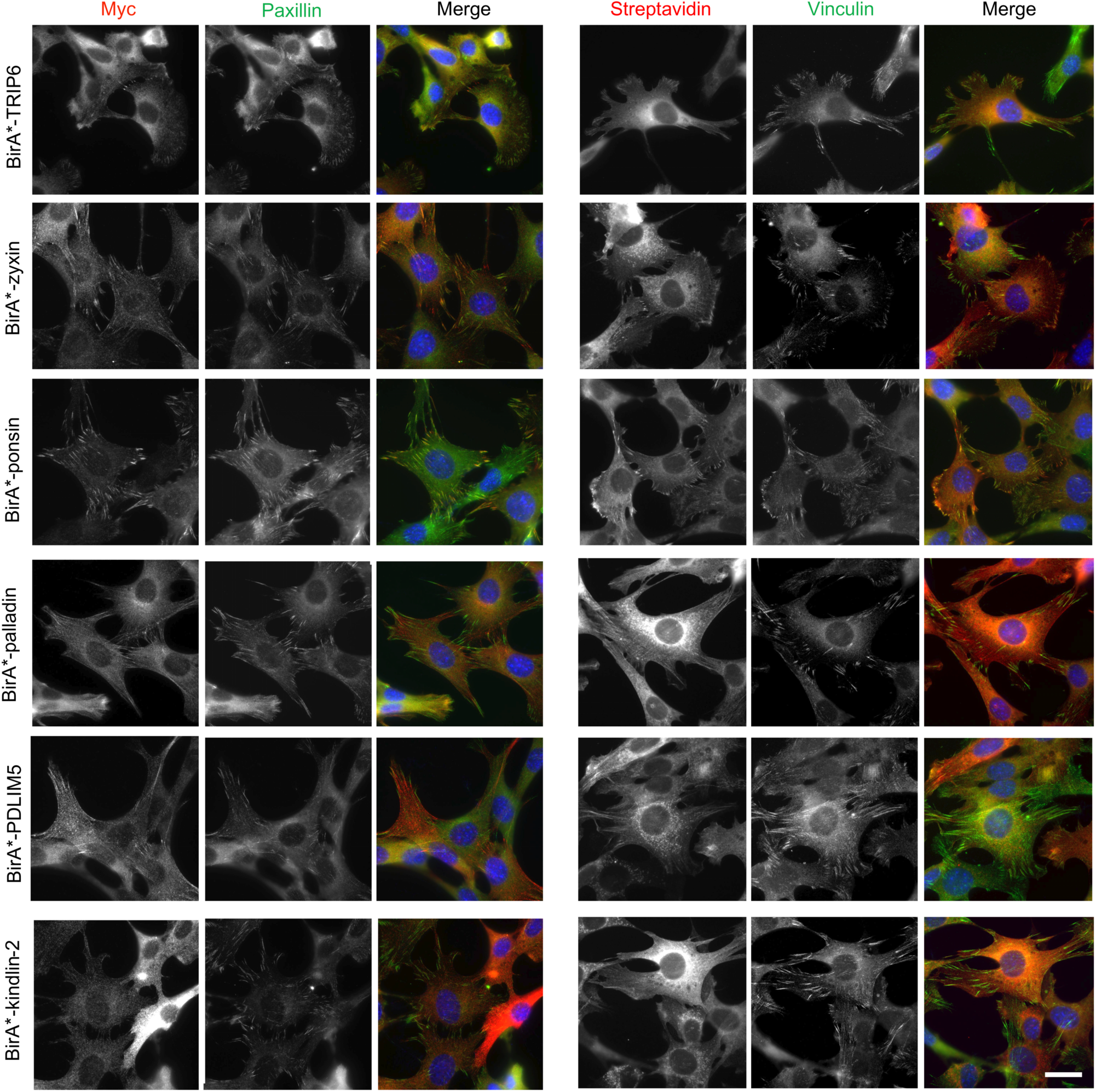
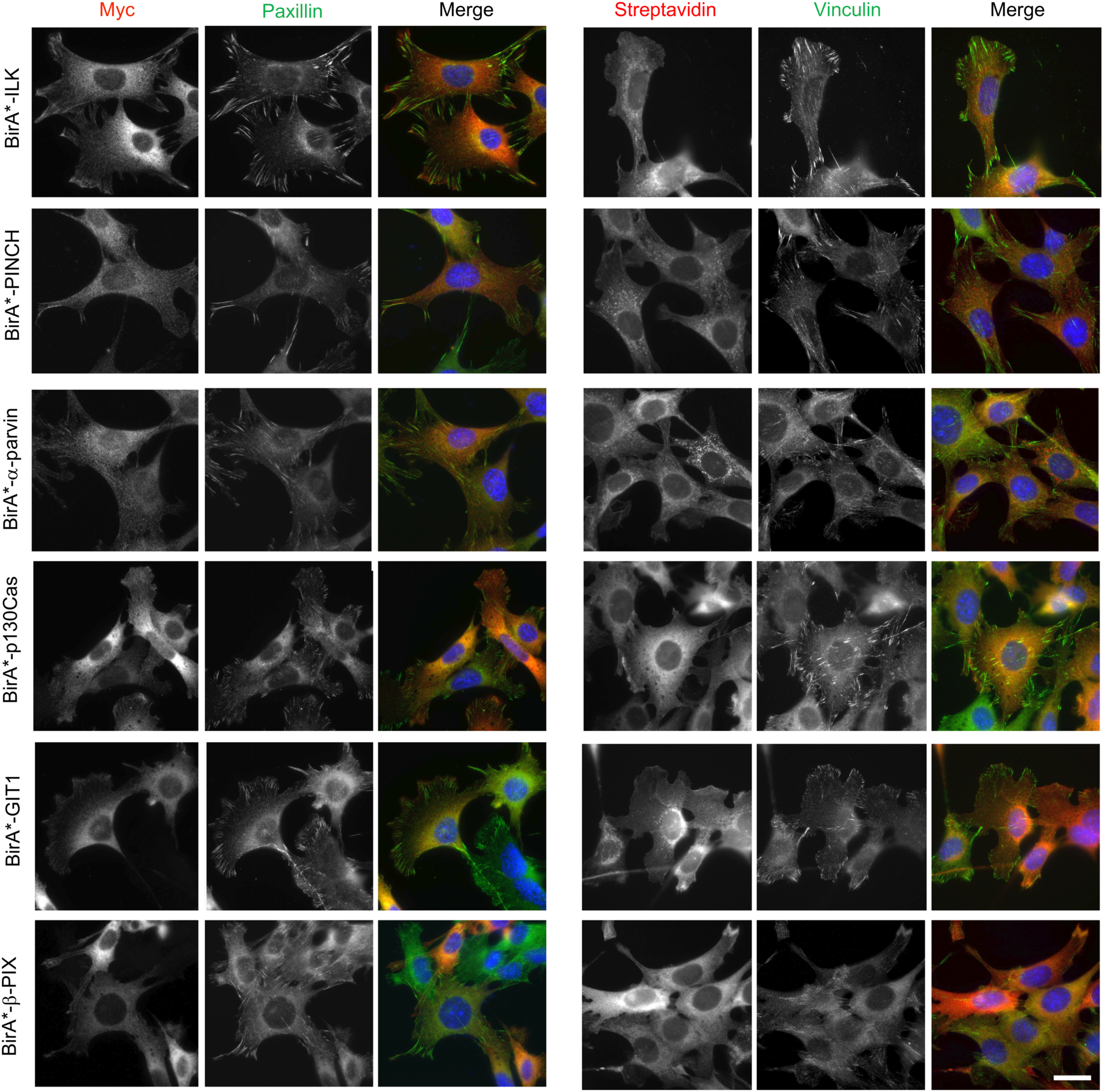
Subcellular localisation of BirA*-tagged adhesome components and biotinylation of proximal proteins. imPSC cells stably expressing BirA* and BirA*-tagged adhesome constructs or untransfected control cells were incubated with biotin for 24 hours before being fixed and stained for myc and paxillin, or vinculin and biotinylated proteins (using fluorophore-conjugated streptavidin). Scale bar: 30 μm.

**Supp. 2.**
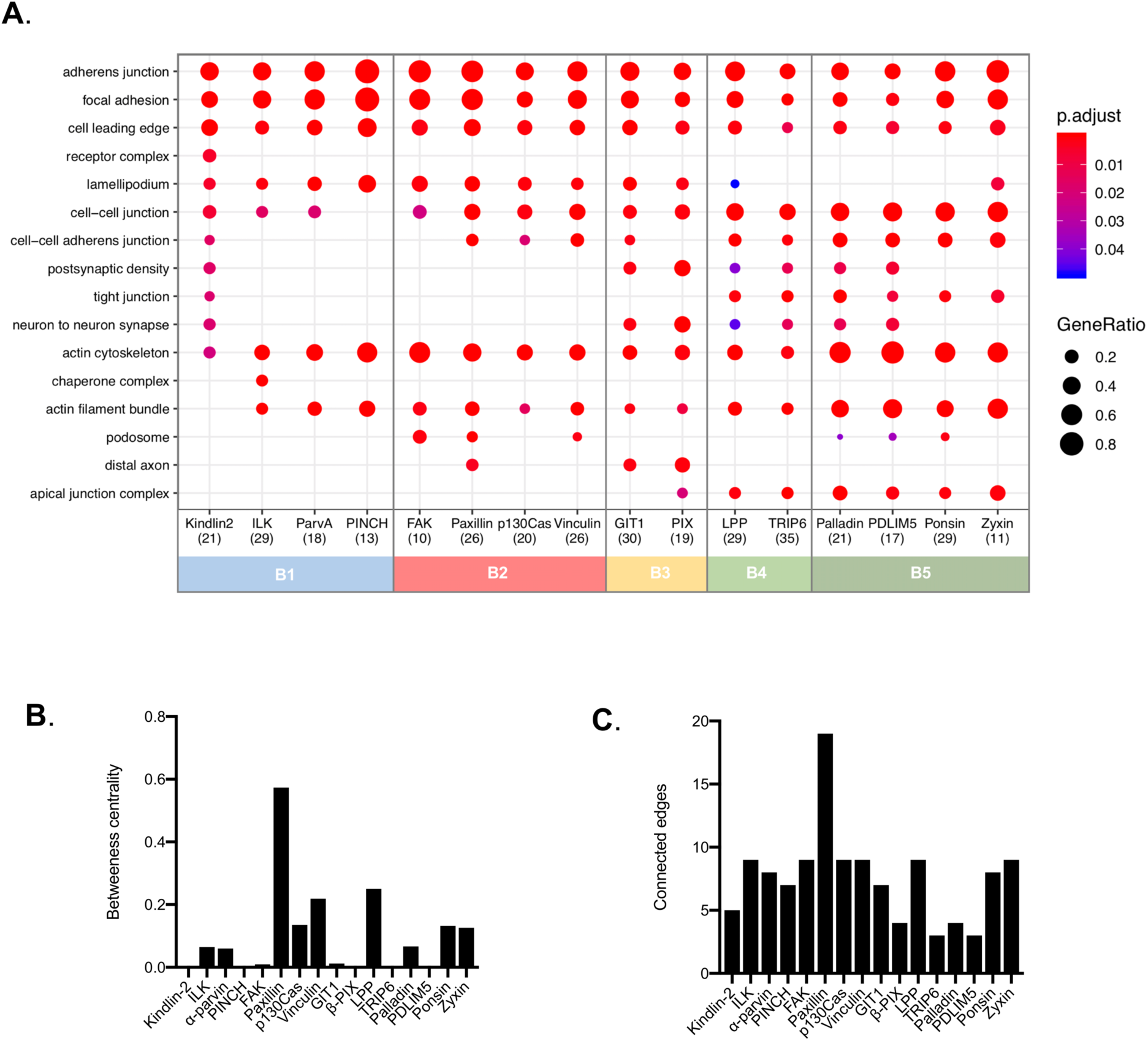
Ontological analysis of adhesome BioID data. **A.** Functional enrichment analysis of proteins identified by each of the 16 BirA*-tagged adhesome proteins (BFDR ≤ 0.05). The top three over-represented GO terms under the ‘cellular component’ category for each bait are listed, and displayed for all baits (if identified) with an adjusted p-value ≤ 0.05. The number of proteins recognised per interactome is shown in brackets. Baits are ordered and colour-coded according to hierarchical clustering as in fig. 2. p.adjust: adjusted p-value. GeneRatio: proportion of total proteins identified in each GO term. **B.** Betweenness centrality of each bait from network analysis of bait-prey interactions between BirA*-tagged baits in fig. 3B. **C.** Number of connected edges of each bait from network analysis of bait-prey interactions between BirA*-tagged baits in fig. 3B.

**Supp. 3.**
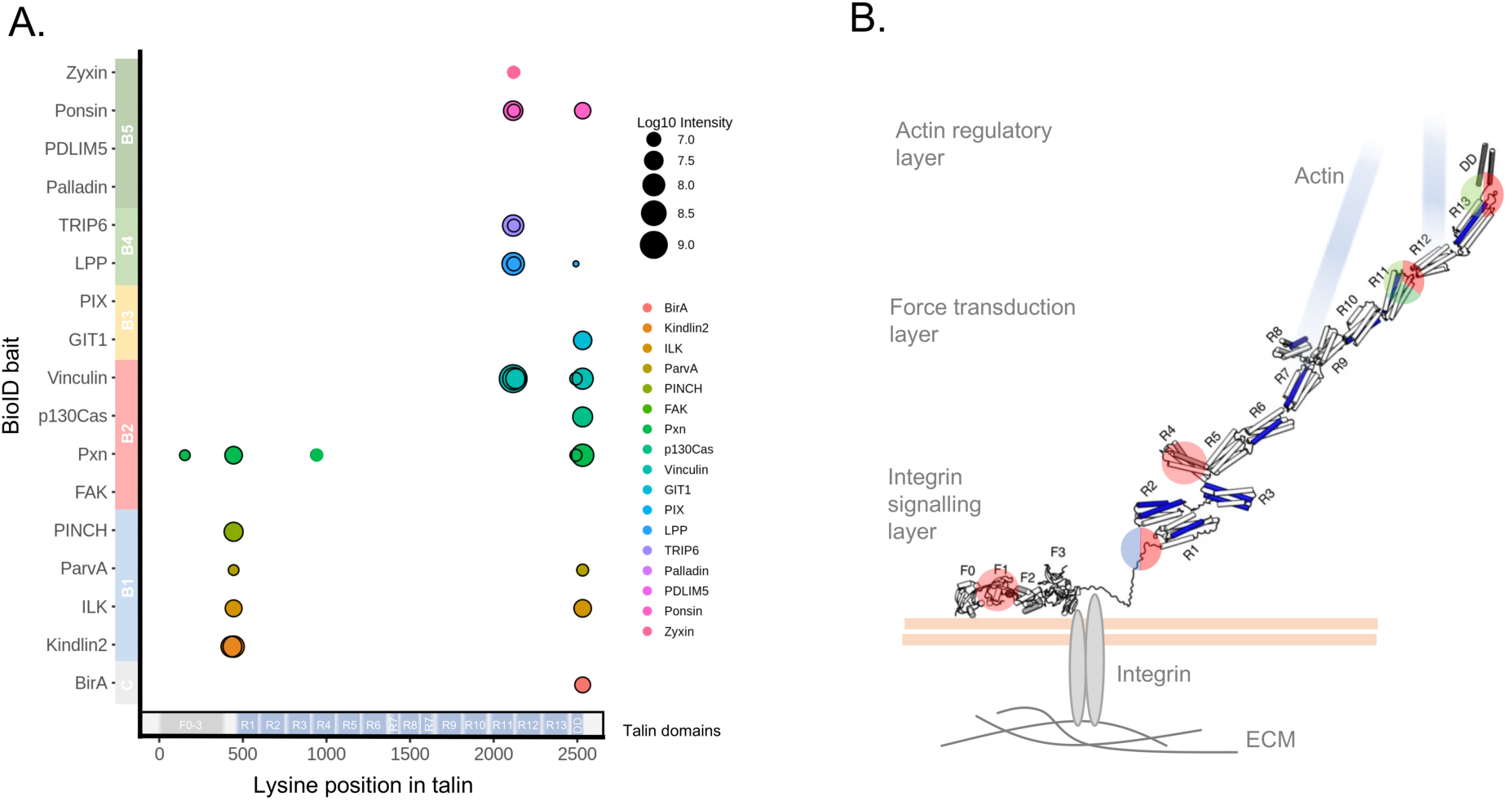
Talin biotinylation by adhesome baits. **A.** Biotinylated lysines from talin identified by each adhesome bait were mapped onto the talin sequence. Black borders indicate biotinylated peptides present in a minimum of two of three repeats. Baits are coloured according to the hierarchical clustering, and the domain structure of talin is indicated at the bottom 35. Note that the most C-terminal biotinylated peptide in the DD domain is also found in BirA* alone and therefore likely to be non-specific. **B.** Schematic of biotinylated lysines mapped onto the tertiary structure of talin in its extended active form in IACs. Vinculin binding sites are shown in blue. The position of biotinylated lysines are highlighted and colour-coded according to the BioID bait clusters in figure 2. Blue, B1; red, B2; light green, B4; dark green, B5. Talin structure adapted from Yao *et al* 102. F0-3, FERM domains 0-3; R1-13, rod-domains 1-13; DD, dimerization domain.

**Supp. 4.**
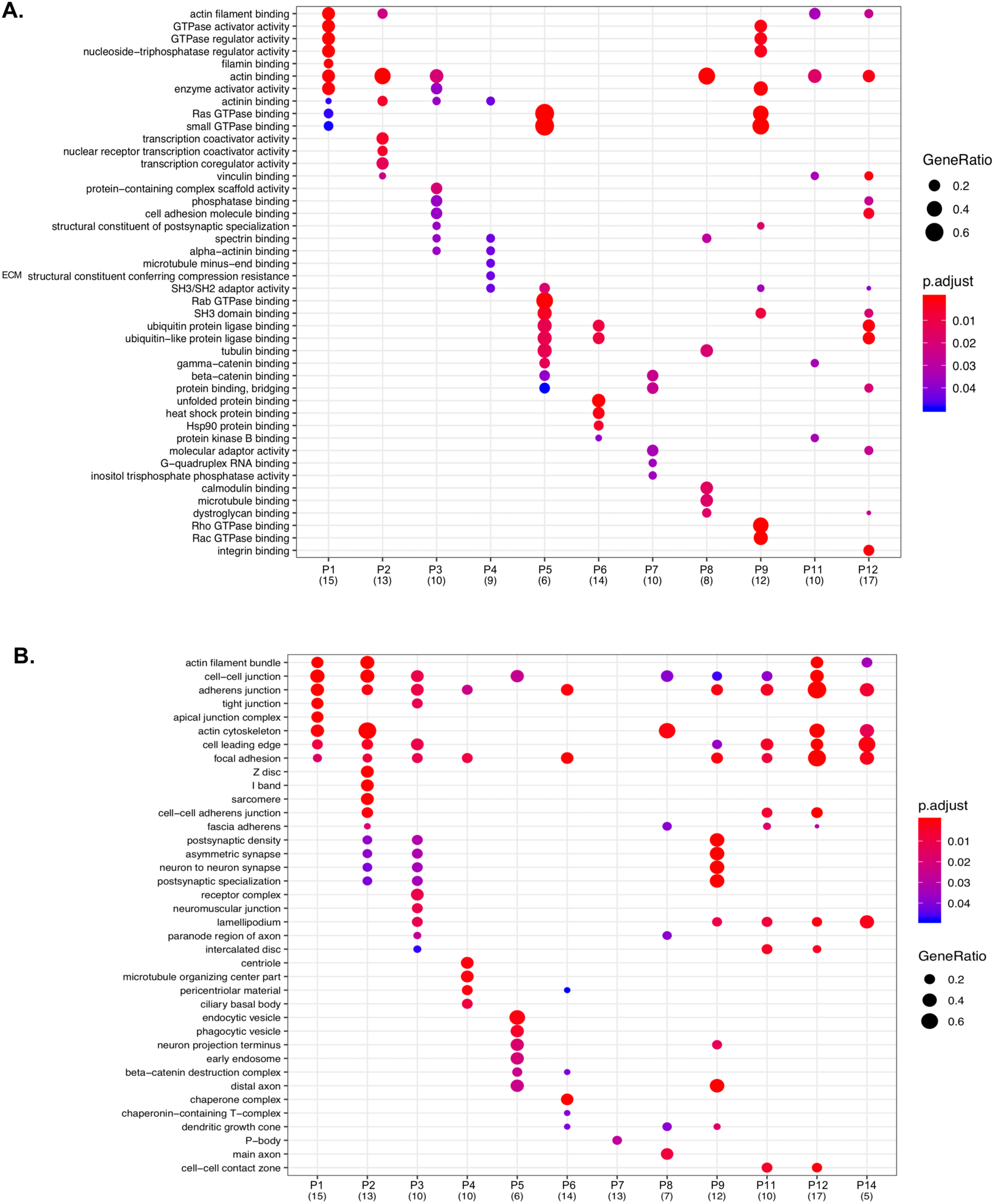
Functional enrichment analysis of prey clusters. GO analysis of the prey clusters identified from hierarchical clustering of the proximity-dependent adhesome (fig. 2) with a minimum of 5 proteins. The top five over-represented GO terms under the ‘molecular function’ **(A)** or ‘cellular component’ **(B)** category for each prey cluster are listed, and displayed for each cluster if identified with an adjusted p-value ≤ 0.05. The number of proteins per prey cluster is shown in brackets. p.adjust: adjusted p value. GeneRatio: proportion of total proteins identified in each GO term.

Supp. table 1. The proximity-dependent adhesome

Supp. table 2. Hierarchical clustering of BioID baits and prey

Supp. table 3: Biotinylation of lysines in talin-1

Supp. table 4: List of primers and primer pairs used to generate BioID constructs

